# When less is more: Enhanced statistical learning of non-adjacent dependencies after disruption of bilateral DLPFC

**DOI:** 10.1101/198515

**Authors:** Géza Gergely Ambrus, Teodóra Vékony, Karolina Janacsek, Anna B. C. Trimborn, Gyula Kovács, Dezso Nemeth

## Abstract

Brain networks related to human learning can interact in cooperative but also competitive ways to optimize performance. The investigation of such interactive processes is rare in research on learning and memory. Previous studies have shown that manipulations reducing the engagement of prefrontal cortical areas could lead to improved statistical learning performance. However, no study has investigated how disruption of the dorsolateral prefrontal cortex (DLPFC) affects the acquisition and consolidation of non-adjacent second-order dependencies. The present study aimed to test the role of the DLPFC, more specifically, the Brodmann 9 area in implicit temporal statistical learning of non-adjacent dependencies. We applied 1 Hz inhibitory transcranial magnetic stimulation or sham stimulation over both the left and right DLPFC intermittently during the learning. The DLPFC-stimulated group showed better performance compared to the sham group after a 24-hour consolidation period. This finding suggests that the disruption of DLPFC during learning induces qualitative changes in the consolidation of non-adjacent statistical regularities. A possible mechanism behind this result is that the stimulation of the DLPFC promotes a shift to model-free learning by weakening the access to model-based processes.

## Introduction

Accumulating evidence supports the existence of interactive learning/memory processes, which can be either cooperative or competitive. Such a competitive relationship is theorized to exist between the instructional/deliberate *model-based* and the incidental/reflexive *model-free* processes underlying learning (Daw, Niv, & Dayan, 2005). Model-based processes refer to the more controlled forms of learning that include the development of complex representations based on testing hypotheses about the environment (Beierholm, Anen, Quartz, & Bossaerts, 2011; Wan Lee, Shimojo, & O’Doherty, 2014). Consequently, internal models are formulated, which may be used both for the already learned and the newly encountered structures (Daw et al., 2005; Haith & Krakauer, 2013). These processes were associated with goal-directed control, processing speed, executive functions, and working memory (e.g., Kurth-Nelson, Bickel, & Redish, 2012; Otto, Raio, Chiang, Phelps, & Daw, 2013; Schad et al., 2014). Model-free processes refer to habit-like, associative forms of learning whereby one extracts predictable structural regularities from the environment without intention or conscious monitoring. Thus, this type of learning is stimulus-driven and typically occurs implicitly. This predictive processing and capacity to detect patterns are crucial for aspects of *statistical learning,* which are involved in the acquisition of cognitive, social, and motor skills and habits (e.g., Kaufman et al., 2010; Lieberman, 2000; Nemeth & Janacsek, 2011). Disruptive stimulation of the dorsolateral prefrontal cortex (DLPFC) was shown to shift the balance of model-free and model-based processes to the benefit of model-free processes during reinforcement learning (Smittenaar, FitzGerald, Romei, Wright, & Dolan, 2013). Congruently, better statistical learning was associated with weaker model-based processes (Janacsek, Fiser, & Nemeth, 2012; Nemeth, Janacsek, Polner, & Kovacs, 2013; Tóth et al., 2017; Virag et al., 2015), supporting the potential competition between the two types of learning processes. Can “less” involvement of the DLPFC be “more” beneficial for cognitive functions driven by model-free processes? Our study aimed to answer this question by directly manipulating the involvement of DLPFC in temporally distributed statistical learning using repetitive transcranial magnetic stimulation (rTMS).

Previous research has proposed that the neural circuitry involving parts of the basal ganglia, notably the dorsolateral striatum, supports model-free learning processes. The circuitry involving medial temporal lobe structures, including the hippocampus, and the areas of the default network (Buckner & DiNicola, 2019; Daw, Gershman, Seymour, Dayan, & Dolan, 2011; Dayan & Berridge, 2014; Janacsek et al., 2012; Packard & Knowlton, 2002; Vikbladh et al., 2019; Wunderlich, Smittenaar, & Dolan, 2012; Yin, Knowlton, & Balleine, 2004) promotes model-based processes. Studies suggest that both processes involve the lateral prefrontal cortical regions that subserve several cognitive functions, such as executive functions, working memory, memory encoding, and access to long-term memory (Baier et al., 2010; Blumenfeld & Ranganath, 2007; Culbreth, Westbrook, Daw, Botvinick, & Barch, 2016; Koechlin & Summerfield, 2007; Lara & Wallis, 2015; McNab & Klingberg, 2008; Otto et al., 2013; Otto, Skatova, Madlon-Kay, & Daw, 2014). The neural basis of picking up probabilistic statistical regularities has been investigated by multiple neuroimaging studies. For example, Simon, Vaidya, Howard, and Howard (2012) used event-related functional magnetic resonance imaging during a probabilistic sequence learning task. The study provided evidence for the role of the hippocampus in the early stage of learning and the caudate in later stages. Using diffusion tensor imaging, Bennett, Madden, Vaidya, Howard, and Howard (2011) found that the integrity of the neural tracts between the DLPFC and the hippocampus, and also between the DLPFC and the caudate nucleus are related to the degree of statistical learning. Stillman et al. (2013) found that a functional connectivity index between the caudate and medial temporal lobe showed a positive correlation with probabilistic statistical learning. Based on their findings, the authors also stressed the potential mediating role of the DLPFC between other, statistical learning-related areas such as the caudate and the medial temporal regions. Thus, we hypothesized that the DLPFC might modulate statistical learning abilities by being a part of the neural circuits, both supporting the model-based and model-free processes.

Employing non-invasive brain stimulation methods, we can reveal relationships between learning and brain areas (and their related networks). To date, only a few studies have investigated the acquisition of temporally distributed deterministic or probabilistic regularities (often termed statistical learning as well) by stimulating the DLPFC. An early rTMS study by Pascual-Leone, Wassermann, Grafman, and Hallett (1996) showed that 5 Hz rTMS over the contralateral DLPFC during a deterministic serial reaction time task (SRTT) impairs online learning. Galea, Albert, Ditye, & Miall (2010) applied inhibitory continuous theta-burst stimulation (cTBS) over the DLPFC following practice on SRTT. After an 8-hour consolidation period, subjects were faster on sequence compared to random elements in the verum, but not in the sham group. Smalle, Panouilleres, Szmalec, and Möttönen (2017) found increased learning on phonological sequences following cTBS over the left DLPFC. Contrary, Savic, Cazzoli, Müri, & Meier (2017) reported null effects of different brain stimulation methods over the DLPFC on a deterministic sequence learning task that requires the use of only one hand. This result suggests that, in some cases, a robust interhemispheric compensation might obscure the effects of the stimulation. More importantly, all the mentioned studies used deterministic sequences with adjacent regularities instead of probabilistic sequences with non-adjacent regularities (Remillard, 2008). In non-adjacent statistical learning, the predictable information is hidden in noise within the stimulus stream. Therefore, the task mimics the acquisition of real-life skills (for example, language learning, Christiansen and Chater, 2015), which occurs under uncertainty, in a noisy environment, more closely (Fiser, Berkes, Orbán, & Lengyel, 2010). The role of the DLPFC in this complex, ecologically valid form of statistical learning remains unclear.

In this study, we go beyond previous findings by disrupting the DLPFC during a non-adjacent statistical learning task using bilateral stimulation. We aimed to reveal whether less involvement of the DLPFC could be more beneficial for the acquisition and consolidation of statistical learning skills. To assess statistical learning, we chose a widely used probabilistic sequence learning task, namely the Alternating Serial Reaction Time (ASRT) task (Figure 1A). This task has been used previously in experimental psychology (Howard & Howard, 1997; Howard et al., 2004; Kóbor, Janacsek, Takács, & Nemeth, 2017; Nemeth et al., 2010; Song, Howard, & Howard, 2007), developmental (Janacsek et al., 2012; Juhasz, Nemeth, & Janacsek, 2019; Nemeth, Janacsek, & Fiser, 2013) as well as neuroimaging studies (Bennett et al., 2011; Stillman et al., 2013). The ASRT task is a four-choice reaction time task in which predetermined stimuli alternate with random elements, creating a probabilistic structure with more probable versus less probable stimulus triplets. Participants can pick up these non-adjacent statistical regularities: over time, performance differences emerge between high- and low-probability triplets without the participants becoming aware of the underlying structure (Howard & Howard 1997; Howard et al. 2004). During the Training/rTMS session, we used 1 Hz repetitive TMS over the DLPFC, which was shown to decrease activity in the targeted brain area (Groiss, Ugawa, Paulus, & Huang, 2012) extending beyond the termination of the stimulation (Robertson, Théoret, & Pascual-Leone, 2003; Walsh & Cowey, 2000). As ASRT requires the use of both hands, bilateral stimulation might be an ideal choice to control for the possible premotor response bias (by bilateral stimulation, high- and low-probability triplets are affected to a similar extent).

**Figure 1.**
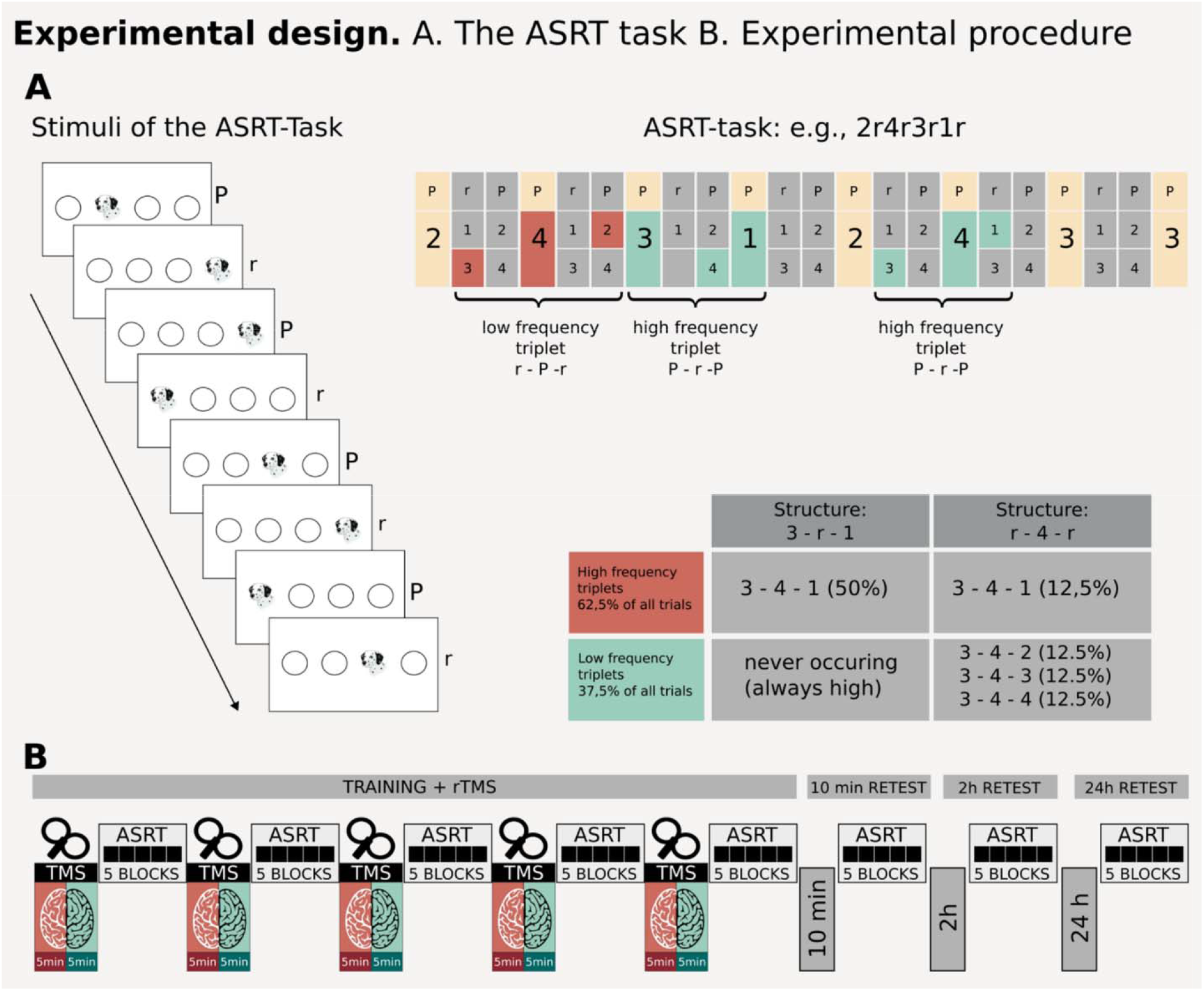
Experimental procedures. (A) Stimuli of the Alternating Serial Reaction Time (ASRT) task. Repeating elements (P – pattern) alternate with random events (r – random). Due to this structure of the sequences, some triplets (i.e., three consecutive events) occur more frequently (high-probability triplets) than others (low-probability triplets). Implicit statistical learning is measured as the RT difference between these two triplet types. (B) Five minutes of 1 Hz rTMS of both DLPFCs was administered before each of the five learning blocks, with the order of the stimulated hemispheres counterbalanced inter-participants. The volunteers performed five ASRT blocks 10 minutes, 2 hours, and 24 hours post-learning, as well.

Moreover, a sequential bilateral TMS protocol was chosen to suppress the compensation of the non-stimulated hemisphere. Retest sessions 10 minutes, 2 hours, and 24 hours after the termination of the training were implemented in the design. Our purpose was to study the effect of TMS on the whole learning process and to differentiate between the immediate and long-lasting results of stimulation and its impact on the offline consolidation of acquired knowledge. We hypothesized that disrupting the DLPFC bilaterally would increase statistical learning performance in the ASRT task.

## Materials and methods

### Participants

Thirty-two participants took part in the experiment. All of them were right-handed; their visual acuities were normal or corrected to normal. None of the participants reported a history of neurological or psychological disorders, drug or alcohol abuse, had metal implants, or were taking regular medication relevant to the study. Written informed consent was acquired from all participants. All participants tolerated the experimental procedures, and none withdrew because of discomfort with TMS. All participants were students of the University of Jena and participated in exchange for partial course credits or monetary compensation. One participant was excluded from the final sample because of poor performance on the Berg Card Sorting Test (% of preservative errors = 31.25%; % of correct responses: 51.56%; the scores were more than three standard deviations above the mean, see group averages in the Experimental Procedure section). Thus, the final sample contained data from 31 participants (16 in the DLPFC Group and 15 in the Sham Group, four males, M_age_ = 22.16 years, SD_age_: ± 3.01). The experiment was conducted in accordance with the guidelines of the Declaration of Helsinki and with the approval of the ethics committee of the University of Jena.

### Alternating Serial Reaction Time Task

Statistical learning was measured using the Alternating Serial Reaction Time (ASRT) Task (Howard et al., 2004; Song et al., 2007). In this task, a stimulus (a dog’s head) appeared in one of four horizontally arranged empty circles on the screen. Participants were instructed to press the corresponding key (Z, C, B, and M on a QWERTY keyboard), as quickly and as accurately as possible (Figure 1A). The buttons Z and C had to be pressed by the middle and index fingers of the left hand. The B and M buttons had to be pressed by the index and middle fingers of the right hand, respectively. The target remained on the screen until the participant pressed the correct button. The response-to-stimulus interval was set to 120 ms. Stimuli were presented in blocks of 85 trials. The first five trials of each block were random elements and for practice purposes only (not analyzed further). After these five practice trials, an eight-element alternating sequence was repeated ten times in a block (e.g., 2*r*4*r*3*r*1*r*, where 1–4 indicate the target locations from left to right, and *r* indicates a randomly selected position out of the four possible ones) (Figure 1A). The predetermined order of the pattern elements remained unknown to the participants. Due to the alternation of random and pattern elements, some runs of three consecutive items (henceforth referred to as triplets) occurred with higher probability than other ones. We refer to these types of stimuli as high-probability and low-probability triplets, respectively. For example, considering the above illustration, 2_**4**, 4_**3**, 3_**1**, and 1_**2** (where “_” indicates the middle element of a triplet) occur with high-probability, because the third element (bold number) could be derived from a (predetermined) pattern, and in some cases, from random items. On the contrary, 1_**3** and 4_**1** occur with less probability because in that case, the third element could only be random. Therefore, the third event of a high-probability triplet is more predictable from the first element than in the case of the low-probability triplets. Accordingly, each of the trials of the ASRT was categorized as either the third element of a high- or a low-probability triplet. Overall, 64 possible triplets can occur in the task, of which 16 are high-probability triplets, each of them occurring in approximately 4% of the trials (62.5% in total). Each of the remaining 48 triplets occurred in around 0.8% of the trials (37.5% in total). Thus, the high-probability triplets occur five times more often than the low-probability triplets.

### Structural MRI and Neuronavigated TMS

Structural MRI scanning was performed using a Siemens Magnetom Trio 3T MRI scanner at the Institute for Diagnostic and Interventional Radiology, University of Jena. High-resolution sagittal T1-weighted images for the 3D head and brain meshes were acquired using a magnetization EPI sequence (MP-RAGE; TR = 2300 ms; TE = 3.03 ms; 1 mm isotropic voxel size). For neuronavigated TMS, the 3D-head and brain models were created from the participants’ individual MRI scans. Coordinates for the DLPFC were taken from a meta-analysis by Cieslik et al. (2013), which corresponds to the dorsal part of the DLPFC (MNI coordinates: x = 37, y = 33, Z = 32). This area was revealed to be involved in executive control and working memory processes. For sham TMS, the coil was oriented perpendicularly, facing away from the skull (Lisanby, Gutman, Luber, Schroeder, & Sackeim, 2001).

TMS stimulation was delivered using a PowerMag 100 Research Stimulator (MES Forschungssysteme GmbH). Neuronavigation was carried out using a PowerMag View (MES Medizintechnik GmbH) Neuronavigation system. Magnetic pulses were delivered at 1 Hz, at 55% maximum stimulator output. A single intensity was used based on previous studies (Figner et al., 2010; Silvanto, Cattaneo, Battelli, & Pascual-Leone, 2008). TMS was applied before each of the five learning blocks, that is, before the first block and in the inter-block intervals (300 pulses, 5 minutes per hemisphere). The order in which the two hemispheres were stimulated was counterbalanced inter-participants.

### Experimental procedures

Participants were seated in a dimly lit room; their heads were fixed using a chinrest, 60 cm viewing distance away from the stimulus presentation monitor. After giving informed consent, the volunteers performed an ASRT practice run to familiarize themselves with the task and the keyboard layout. In the Training/rTMS session, the participants received 1 Hz rTMS over both left and right hemispheres sequentially (5 minutes, 300 TMS pulses for each hemisphere; thus, 5 minutes for the left and after that the right, in a counterbalanced order inter-participants), then performed five blocks of the ASRT task, lasting approximately 5 minutes. This procedure was repeated five times. Therefore, a total of 25 blocks of ASRT were completed, which provides enough trials for learning to manifest as shown by previous studies (Janacsek et al., 2012; Nemeth et al., 2010; Vékony et al., 2019). The order in which the two hemispheres were stimulated was assigned randomly, remained the same for each participant. To test the role of the DLPFC in statistical learning in both the acquisition and the consolidation, we tested ASRT performance multiple times after the stimulation as well. In the retest sessions, the participants performed five blocks of the ASRT task 10 minutes, 2 hours, and 24 hours after the completion of the Training/rTMS session (Figure 1B). The exact retest times followed previous literature (Janacsek, Ambrus, Paulus, Antal, & Nemeth, 2015). The 10 minutes retest session aimed to test the immediate aftereffect of TMS *without* preceding stimulation. With the help of the 2 hours retest session, we could test the long-term effects of the stimulation. The performance after 24 hours was thought to provide information about the impact of the stimulation on consolidation.

To ensure that the two experimental groups did not differ in executive function performance, we administered the short-form of the Berg Card Sorting Test (Fox, Mueller, Gray, Raber, & Piper, 2013) and the Counting Span test (Case, Kurland, & Goldberg, 1982; Conway et al., 2005; Engle, Laughlin, Tuholski, & Conway, 1999) after the completion of the ASRT in the last session. We observed no significant differences in performance between the two experimental groups (Berg Card Sorting Test, percent correct responses: DLPFC: M = 81.06, SD = 5.90; Sham: M = 80.42, SD = 7.76, *p* = .80; percent perseverative errors: DLPFC: M = 10.16, SD = 4.56; Sham: M = 11.67, SD = 4.83, *p* = .38; percent non-perseverative errors: DLPFC: M = 8.79, SD = 5.59; Sham: M = 7.92, SD = 5.06, *p* = .65; Counting span, mean of three runs: DLPFC: M = 4.06, SD = 1.11, Sham: M = 3.62, SD = 0.85, *p* = .23).

As a part of the post-experimental debriefing, the participants filled out a questionnaire assessing their levels of discomfort, tiredness, and perceived task difficulty, measured on a tenpoint scale.

The Training/rTMS session (with informed consent) lasted approximately 2 hours, the 10 minutes and 2 hours retest sessions lasted 5 minutes each, and the 24 hours retest session (with the control tasks) and debriefing lasted approximately 30 minutes.

### Statistical analysis

Only correct responses were considered for the ASRT analysis, and stimulus repetitions (e.g., 333, 444) and trills (e.g., 313, 121) were also excluded (Howard & Howard, 1997; Howard et al., 2004). Trials with reaction times (RTs) more than 2.5 standard deviations above or below the mean of the given epoch were eliminated (separately for each participant). After that, we calculated the mean RTs for high- and low-probability triplets separately. Implicit statistical learning was assessed using a triplet-learning index, calculated by subtracting the RTs for low-probability triplets from those for high-probability triplets. To control for the non-specific effects of time and the possible individual differences in RTs, we calculated a percentage learning index as follows: [(RTs for low-probability triplets - RTs for high-probability triplets)/RTs for low-probability triplets)]. A higher learning index thus means faster responses for high-than for low-probability triplets, that is, better statistical learning performance.

The learning index was calculated for each session of the experiment for each participant. We conducted a 2 (Group: DLPFC Stimulation vs. Sham Stimulation) × 4 (Session: Training/rTMS session vs.10 minutes retest session vs. 2 hours retest session vs. 24 hours retest session) mixed-design ANOVA to compare the statistical learning performance between the two stimulation groups throughout the experiment. Greenhouse-Geisser corrections were applied where necessary. Significant main effects and interactions were further analyzed using Bonferroni-corrected post-hoc comparisons.

As we aimed to investigate the effects of rTMS on both hemispheres, we tested whether the observed effect was due to the stimulation of the hemisphere just before the particular ASRT block. Thus, we conducted a 2 (Order: Right Start vs. Left Start) × 4 (Session) ANOVA on the learning index to ascertain if our results were influenced by the stimulation order.

We also performed an analysis of the ASRT performance across the four sessions with the raw RTs for high- and low-probability triplets as dependent variables. To see how the initial learning was affected by the stimulation, we ran an analysis of the learning indices of the five epochs of the Training/rTMS session. We also compared the learning indices of the three retest sessions to the last epoch of the Training/rTMS session (see details of these analyses and results in the Supplementary Materials).

All analyses were two-tailed and were conducted with a significance level of *p* < .05.

## Results

The ANOVA conducted on the learning indices across sessions revealed a significant main effect of the experimental Session, *F*_3, 87_ = 7.11, *p* < .001, η_*p*_^2^ = .20. The pairwise comparisons of this main effect indicate an overall increase in performance in all three retest sessions when compared to the Training/rTMS session (Training/rTMS session vs. 10 minutes retest session: *p* = .001, 2 hours retest session: *p* = .001, 24 hours retest session: *p* = .004). There was no main effect of Group, *F*_1, 29_ = 0.61, *p* = .44, η_*p*_^2^ = .02, but the interaction between the experimental Session and Group was shown to be significant, F_3, 87_ = 3.96,*p* = .01, η_*p*_^2^ = .12. In the Sham Group, pairwise comparisons revealed an increased performance at the 10 minutes retest session compared to the Training/rTMS session (*p* = .002). Performance in the DLPFC Group also increased with time compared to the Training/rTMS session (2 hours retest session: *p* = .01), and contrary to the Sham Group, the difference remained significant at the 24 hours retest session (*p* = .001) (Figure 2A). The comparisons of the learning indices between the two groups revealed that the Sham and DLPFC Groups showed similar learning indices during the Training/rTMS, 10 minutes, and 2 hours sessions (all *p* > .25). However, the DLPFC Group showed a significantly greater learning index than the Sham Group at the 24 hours retest session (*p* = .03) (Figure 2B). The stimulation seemed to affect the RTs for the low-probability triplets primarily; see details in the first section of the Supplementary Materials.

**Figure 2.**
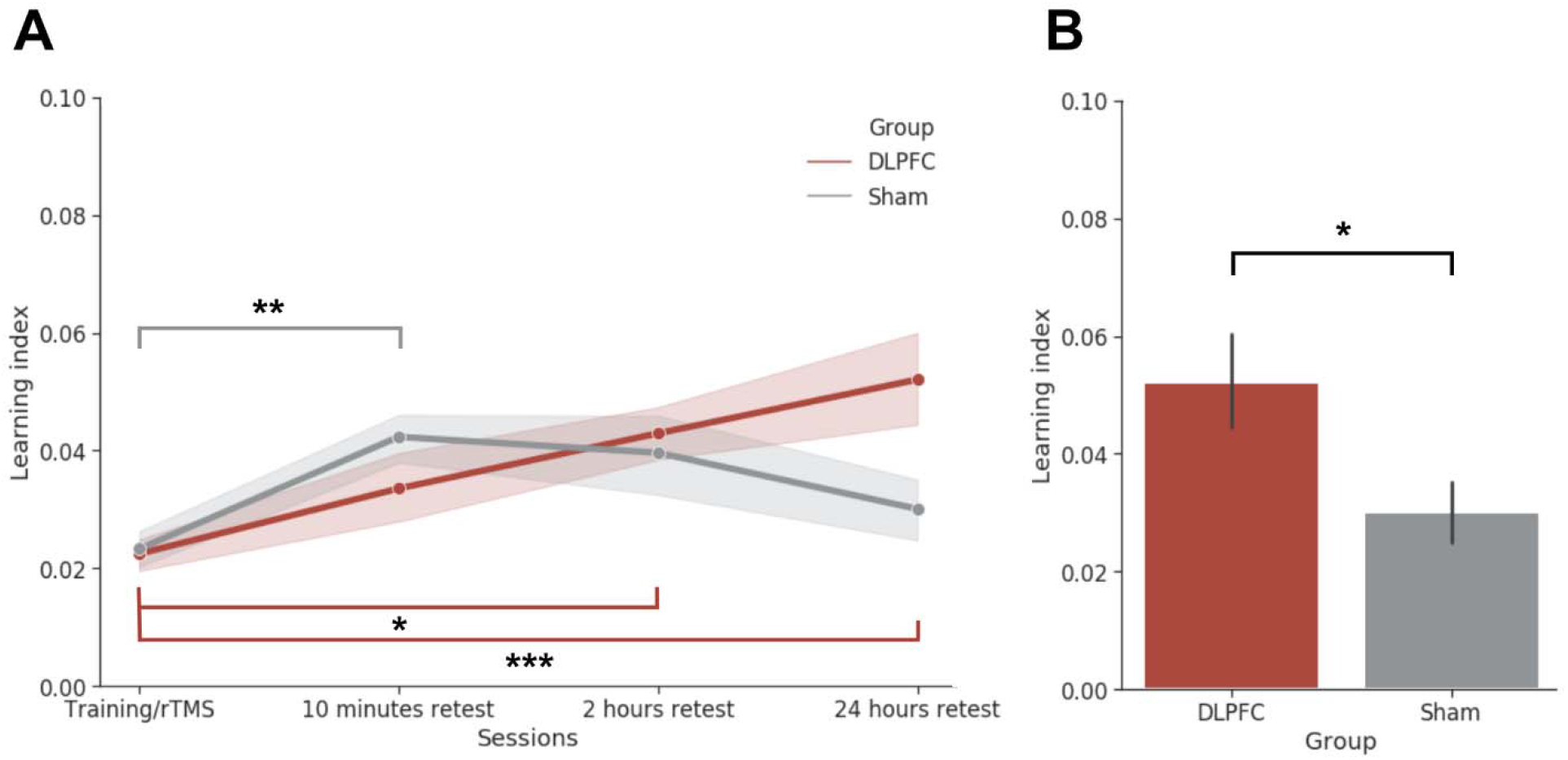
(A) The learning index in the two experimental groups along the course of the experiment and (B) the comparison of the triplet learning indices between groups in the last session. Compared to the Training/rTMS session, we observed an increase in performance in the active DLPFC Group. However, the performance in the Sham Group was only significantly better in the 10 minutes retest session (A). After 24 hours, the performance of the active DLPFC Group was statistically better that of the Sham Group (B). The error bars denote the SEM. *: *p* < .05, **: *p* < .01, ***: *p* = .001.

The observed group differences are unlikely to be due to a general effect of the stimulation on arousal level because the general reaction time (Session × Group interaction: *F*_3,87_ = 0.39 *p* = .76, η^2^_*p*_ = .01) and response accuracy (Session × Group interaction: *F*_3,87_ = 0.38, *p* = .77, *η*^2^_*p*_ = .01) was not statistically different between the two groups. Furthermore, the level of discomfort (*p* = .57), tiredness (*p* = .83), perceived task difficulty (*p* = .24), assessed as a part of the post-experiment debriefing, was also not different between the DLPFC and the Sham Group (see Methods).

Moreover, the order of the hemispheres stimulated, assessed by the two-way interaction of a 2 (Order: Right Start vs. Left Start) × 4 (Session) mixed-design ANOVA, did not affect statistical learning performance, *F*_3,42_ = 0.15, *p* = .93, η^2^_*p*_ = .01.

## Discussion

To date, only a few studies have investigated the role of the DLPFC in statistical learning (Galea et al., 2010; Pascual-Leone et al., 1996; Savic et al., 2017; Smalle et al., 2017), while none of them tested its role in the acquisition and consolidation of non-adjacent dependencies. Here, we aimed to fill this gap by administering bilateral rTMS over the DLPFCs during temporal statistical learning of non-adjacent dependencies. We went beyond previous studies in three aspects. First, instead of deterministic sequences, we tested the effect of rTMS on the acquisition of non-adjacent probabilities. Second, we applied bilateral stimulation to control for a possible interhemispheric compensation of the non-stimulated hemisphere. Third, we tested the effect of the TMS protocol used throughout the training session as well as 10 minutes, 2 hours, and 24 hours post-training. Our results show that the bilateral disruption of the DLPFCs during the training session has a beneficial effect on the statistical learning performance after 24 hours. Therefore, we suggest that DLPFCs play a role in non-linguistic statistical learning processes. As predicted, our findings are in line with the competition model that posits an antagonistic relationship between model-based and model-free learning processes (Daw et al., 2011; Janacsek et al., 2012; Nemeth et al., 2013; Smittenaar et al., 2013; Virag et al., 2015).

In agreement with the computational framework of model-free and model-based processes, previous research further demonstrated that cognitive functions that are mainly determined by these two types of processes have an inverse relationship on the behavioral level. For instance, Virag et al. (2015) showed a negative correlation between working memory/executive functions and implicit statistical learning. Filoteo et al. (2010) found that implicit category learning improved with the addition of a secondary working memory task, that is, with the reduction of the contribution of model-based learning processes (however, for a critical re-evaluation of this study see Newell, Moore, Wills, & Milton, 2013). Nemeth et al. (2013) found increased statistical learning performance in hypnosis compared to an alert, awake state, by possibly reducing long-range brain connectivity (Fingelkurts, Fingelkurts, Kallio, & Revonsuo, 2007; Oakley & Halligan, 2009). Additionally, several developmental studies found that children perform better than adults on non-adjacent statistical learning tasks (Janacsek et al., 2012; Juhasz et al., 2019; Zwart, Vissers, Kessels, & Maes, 2019). Supporting the competition model, the degree of learning decreases with the onset of adolescence, coinciding with the maturation of the dorsolateral regions of the prefrontal cortex (Bunge & Zelazo, 2006; Gogtay et al., 2004; Kadosh, Heathcote, & Lau, 2014; Thompson et al., 2001). Besides the abovementioned behavioral studies, Tóth et al. (2017) found evidence for this inverse relationship at the level of neural oscillations. Namely, they detected increased statistical learning associated with weaker fronto-parietal connectivity in theta frequency, a band that plays a crucial role in memory access (Düzel, Penny, & Burgess, 2010) and also in sentence processing and working memory (Beese, Meyer, Vassileiou, & Friederici, 2017). These previous studies revealed only indirect, correlational relationships; however, our recent findings yield evidence for the role of the DLPFC in non-adjacent statistical learning.

To date, only four studies have investigated the role of the DLPFC on temporally distributed deterministic or probabilistic regularities using TMS protocols. Intending to disrupt cortical processing by 5 Hz rTMS during deterministic SRTT, Pascual-Leone et al. (1996) found that stimulation over the contralateral DLPFC impaired online learning. It should be noted that since the publication of this study, 5 Hz rTMS has been found to induce excitatory effects on cortical excitability (Matsunaga et al., 2005; Peinemann et al., 2004); thus, the performance decrease in this study might be better explained by the facilitation of DLPFC functions, in line with the competition account. To test the role of the DLPFC in the consolidation of sequential knowledge, Galea et al. (2010) applied disruptive cTBS after the execution of deterministic SRTT. They found an offline improvement following the inhibition of the right but not the left DLPFC, which they explained by interference between declarative and procedural consolidation processes. However, based on the results of Galea et al., we cannot decide whether a disrupted DLPFC during learning can affect the initial learning. Smalle et al. (2017) went beyond Galea et al. (2010) by applying disruptive stimulation over the left DLPFC prior to learning; increased learning on phonological sequences was found in the Hebb repetition paradigm. A follow-up analysis showed a negative correlation between learning performance and executive functions. To control for the possible compensation of the non-stimulated hemisphere, Savic et al. (2017) tested the effect of brain stimulation over the DLPFC on a deterministic sequence learning task, and no stimulation effect was found over either hemisphere. Taken together, most of the previous TMS studies (except Savic et al. 2017) point in the same direction: facilitating stimulation of the DLPFC hinders, while inhibitory stimulation improves the learning of new sequences, patterns, or statistical regularities.

Our results open up a new theoretical perspective in interpreting the role of the DLPFC in statistical learning. The DLPFC might have a role in model-based processes, such as in accessing the existing models or long-term memory representations, which might be “harmful” when learning of new patterns is required. This idea is supported by results showing that stronger executive functions that substantially involve the activation of the DLPFC might be associated with weaker statistical learning (Janacsek et al., 2012; Smalle et al., 2017; Virag et al., 2015). Moreover, already built models of the statistical regularities seem to hinder the adaptation to changes in those statistical regularities (Kóbor, Horváth, Kardos, Nemeth, & Janacsek, 2019). If access to these model-based processes is limited, then the learning process has to shift towards a model-free approach, which could lead to enhanced learning of entirely new patterns. This framework explains not only the results of the previously mentioned TMS studies on statistical learning well, but also our findings on the consolidation. Namely, as we disrupted the DLPFC, we found better performance after the 24-hour consolidation period. When the DLPFC is fully functioning, the model-free processes extract the statistical information from the stimulus stream, and the DLPFC-mediated model-based processes contaminate these statistics with top-down information. In the offline period, this mixed information consolidates (Figure 3). However, the stimulation of the DLPFC possibly interrupts the top-down information flow and its mixture with the data-driven extraction of pure statistical information. This pure statistical information consolidates, which is optimal when the brain faces the challenge of learning entirely new regularities. At the level of implementation, we suggest that these mechanisms might be realized by modulating the activation in the prefrontal-hippocampal circuitry. It was shown that the neural tracts between the DLPFC and the hippocampus are related to the degree of statistical learning (Stillman et al., 2013). For example, Ross, Brown, and Stern (2009) also showed the role of the hippocampus in the retrieval of learned sequences. A competitive relationship was also described between the hippocampus and striatum during sequence learning, whereby activity in the hippocampus decreases in parallel to an increase in the striatal area (Albouy et al., 2015, 2008). These competitive patterns were linked to performance gains after a consolidation period (Albouy, King, Maquet, & Doyon, 2013). We hypothesize that the DLPFC might play a switching role in similar scenarios (Smittenaar et al., 2013; Stillman et al., 2013). By its disruption, the advantage in the competition changes in favor of the striatal areas. Thus, it can lead to a better consolidation of the statistical regularities because they are more involved in model-free processes.

**Figure 3.**
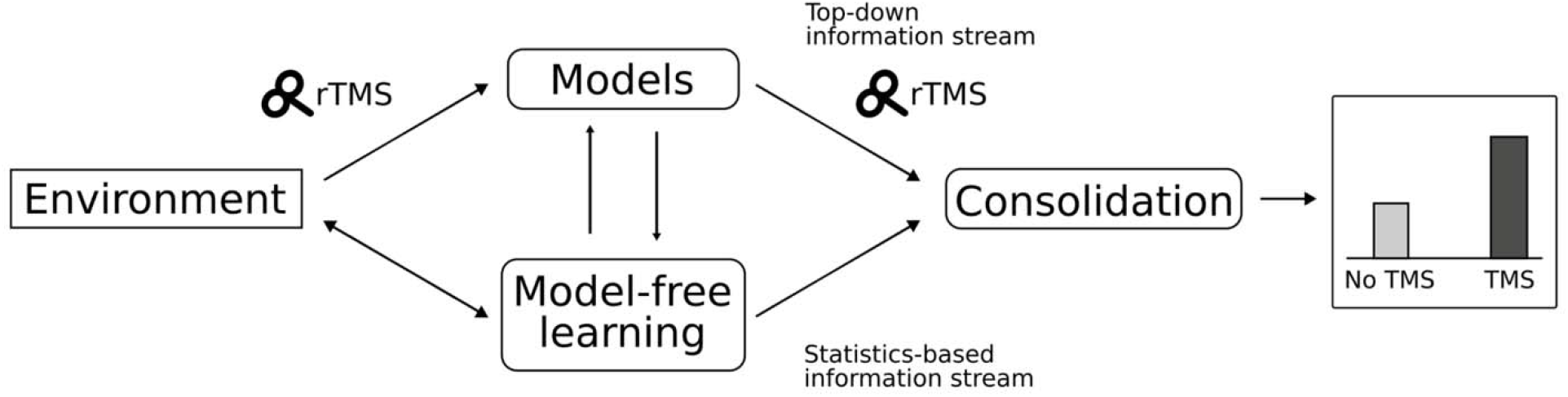
Schematic illustration of the interpretation of our results. Internal models strongly modulate the interpretations of the observed statistics of the input. This helps in extracting complex relations but relatively impairs the detection and learning of raw probabilities. rTMS disrupts the involvement of these internal models, leading to a better consolidation of the newly detected non-adjacent dependencies.

We found an effect of stimulation only after a 24-hour consolidation period. Therefore, a major difference between our results and previous TMS studies is that they found immediate effects of stimulation (Smalle et al. 2017; Pascual-Leone et al., 1996). This difference may be explained by the differences between deterministic and probabilistic statistical learning tasks. In the ASRT task, the relevant information is hidden in the noise; therefore, the model-building process might be slower, and model-free learning might dominate the course of learning through a more extended period. Thus, if stimulation takes place before or during learning, the advantageous effects of reduced model-based processes might appear only at a later time point compared to deterministic tasks. Nevertheless, across-study differences in the stimulation intensity, the number of pulses, or the timing of the stimulation relative to the training session could also contribute to the inconsistencies in the observed aftereffects (Klomjai, Katz, & Lackmy-Vallée, 2015; Thut & Pascual-Leone, 2010). Systematic investigations of the role of the DLPFC in deterministic and probabilistic learning as well as the effects of different stimulation parameters on the learning process, will be required to clarify the inconsistencies in the outcome of stimulation.

We believe that our findings provide a broader theoretical perspective to the process-level understanding of statistical learning. In this view, the fine regulation of shifting between modelbased and model-free processes during learning and consolidation determines the quality of the acquired statistical knowledge. This theoretical framework is supported by evidence using linguistic stimuli (Smalle et al. 2017) and, through the present study, by perceptual-motor stimuli. Based on these findings, we speculate that this phenomenon is generalizable and we hypothesize its existence in most statistical learning situations and tasks. Despite studies investigating the connection between general cognitive functions (such as working memory, attention, executive functions) and statistical learning (Conway, 2020; Frost, Armstrong, & Christiansen, 2019; Janacsek & Nemeth, 2013, 2014), an in-depth theory about their relationship is still missing. The competition framework can open up new research lines to discover the dynamic interactions between general cognitive functions and statistical learning. Future studies should test the competition framework in different statistical learning tasks as well by manipulating the DLPFC.

Our perceptual-motor task with non-adjacent regularities may share similarities with language processes. For example, Nemeth et al. (2011) revealed a relationship between sentence processing and the perceptual-motor statistical learning task used in our study. Here, we found DLPFC-involvement with a non-linguistic perceptual-motor task, and the aftereffects of DLPFC stimulation pointed in the same direction as it was found for a language-related sequence learning task (Smalle et al., 2017). If we broaden the view towards a developmental aspect, language learning and the acquisition of non-adjacent statistical learning appear to share their developmental characteristics: children seem to show better performance both in language learning (Goldowsky & Newport, 1993; Newport, 1990) and perceptual-motor non-adjacent statistical learning (Janacsek et al., 2012; Zwart et al., 2019). Based on the late maturation of the DLPFC (Bunge & Zelazo, 2006; Gogtay et al., 2004; Kadosh et al., 2014; Thompson et al., 2001), we can speculate that the mechanism behind the children’s superiority in these two skills may be related to the effect of TMS over the DLPFC: not having built rigid models about our environment may help liberate our model-free approaches to support the learning of new skills.

This notion is also in line with the finding that the model-based strategy is absent in childhood and gradually strengthens during adolescence up to adulthood (Decker, Otto, Daw, & Hartley, 2016). Future studies directly examining the connection between model-free and model-based processes in language tasks, ideally from a developmental aspect, should be conducted.

Several studies have used non-invasive brain stimulation over Broca’s area to investigate adjacent and non-adjacent dependencies in artificial grammar learning tasks (De Vries et al., 2010; Uddén et al., 2008; Uddén, Ingvar, Hagoort, & Petersson, 2017). Using the same 1 Hz TMS stimulation protocol, they found weaker non-adjacent but better adjacent learning. Therefore, we suggest that the ventral areas (e.g., Broca’s area) might have a different role in the acquisition of non-adjacent dependencies than the dorsal part of the lateral frontal cortex (e.g., Brodmann 9 area, targeted in our study). Yet in natural language, multiple simultaneous non-adjacent dependencies are present (De Vries, Christiansen, & Petersson, 2011), which makes the comparison between the effects on second-order dependencies used in the present study and language learning more complicated. Further research is warranted to reveal the different roles of the dorsal vs. ventral parts of the lateral frontal areas in linguistic learning processes and non-linguistic statistical learning.

Previous studies revealed a possible methodological issue: the interhemispheric compensation might obscure the effects following unilateral stimulation. This effect might have played a role in the negative results of Savic et al. (2017). Galea et al. (2010) found the left hemisphere advantage, but it does not indicate that the activation of the right hemisphere cannot interfere with the results. TMS studies proved that lateralization does not necessarily suggest that the function is eliminated from the other hemisphere, even in the case of language processing (e.g., Hartwigsen et al., 2010) or working memory (e.g., Mottaghy, Döring, Müller-Gärtner, Töpper, & Krause, 2002; Vékony et al., 2018). These results indicate that even if the dominating hemisphere is stimulated, the other can have confounding effects on the results. Thus, we eliminated this possible confounding factor using bilateral brain stimulation to disrupt the involvement of both DLPFCs during learning. Future studies could benefit from using both unilateral and bilateral stimulation in one experimental design to get a holistic picture of the role of the DLPFC in statistical learning.

Finally, there are some limitations to our study. First, we only assessed the level of TMS discomfort after the 24 hours retest session. Thus, the relatively long elapsed time between the stimulation and assessment could have influenced the precision of the subjects’ ratings. The reason for this was that we did not intend to draw the participants’ attention to the perceived (lack of) TMS discomfort. It would have created a belief in the participants about their allocation to the DLPFC or Sham Group, which could have biased our results unnecessarily. Another limitation of our study is that we chose sham stimulation as a control condition: participants were stimulated with a perpendicularly oriented TMS coil. Using such sham control conditions, one can test whether the stimulation of the target area modifies specific processes. However, the regional specificity (i.e., whether stimulation over other regions would not lead to similar changes in performance) can be claimed only with an active control condition (Duecker & Sack, 2015). In our study, the average RTs and accuracy were not altered by the stimulation, suggesting that our results are not due to the modulation of general arousal or attention (Kosinski, 2008), but instead due to the involvement of the targeted DLPFC area in statistical learning itself. Notably, active control stimulation instead of sham stimulation, as suggested above, may also not be optimal as it does not control for placebo effects (Duecker & Sack, 2015). Therefore, in future studies, it would be beneficial to utilize both types of controls within the same experimental design to reveal the regional specificity of the DLPFC for boosting statistical learning.

To sum up, we observed that the bilateral disruption of the DLPFCs during the training session had a beneficial effect on non-adjacent statistical learning performance that was observable after a 24-hour offline period. Our findings are significant in three aspects. First, this finding provides mechanistic level evidence for the models positing an antagonistic relationship between the model-based and model-free processes. Second, from a methodological viewpoint, previous investigations using external non-invasive brain stimulation methods stimulated only one hemisphere at a time. Therefore, the taking-over of the lost function by the contralateral hemisphere cannot be ruled out in earlier studies (Janacsek et al., 2015; Savic et al., 2017). Here, we showed that the sequential application of 1 Hz rTMS before learning blocks over both hemispheres establishes and sustains the inhibitory effect. The finding that no effect of stimulation order was observed supports the viability and practicality of this approach. It may form the basis for future research requiring bi-hemispherical/multi-site intervention. Third, and most importantly, our results raise a new possible theoretical framework explaining the role of the DLPFC in statistical learning and consolidation processes. Our findings shed light on the importance of exploring the possible interactive mechanisms underlying learning. This approach can help us more deeply understand the exact mechanism of skill acquisition and consolidation, and create a bridge between the research fields of general cognitive functions and statistical learning.

## Acknowledgments

This work was supported by a Deutsche Forschungsgemeinschaft Grant (KO3918/5-1, PI: G.K.). This research was supported by the National Brain Research Program (project 2017-1.2.1-NKP-2017-00002); Hungarian Scientific Research Fund (NKFIH-OTKA K 128016, PI: D. N., NKFIH-OTKA PD 124148, PI: K.J.); Janos Bolyai Research Fellowship of the Hungarian Academy of Sciences (to K. J.); IDEXLYON Fellowship of the University of Lyon as part of the Programme Investissements d’Avenir (ANR-16-IDEX-0005) (to D.N). The authors would like to thank Mareike Grotheer and Catarina Amado for performing the MRI scans, Maria Dotzer, Fabienne Windel, and Rebecca Mayer for their help in recruiting the participants.

