## Supplementary Materials for "When less is more: Enhanced statistical learning of non-adjacent dependencies after disruption of bilateral DLPFC"

**Conflict of interest:** The authors report no conflict of interest.

**Analysis of performance for high- and low-probability triplets**

A mixed-design ANOVA with the within-subject factors of Session (Training/rTMS session vs. 10 minutes retest session vs. 2 hours retest session vs. 24 hours retest session) and Triplet (high- vs. low-probability) and with the between-subject factor of Group (DLPFC Stimulation vs. Sham Stimulation) was performed. The main effect of Session was significant, *F*_(3,87)_ = 37.87, *p* < .001, η*_p_^2^* = .57. Note that as here the raw reaction times (RTs) were used as dependent variables, the main effect of Session indicates a change in the average RTs throughout the four sessions. The subsequent post-hoc analysis showed that the RTs in the Training/rTMS session were significantly larger than in the 10 minutes retest session (*p* < .001). In the 2 hours retest session, RTs were larger than in the 10 minutes retest session (*p* < .001). However, for the 24 hours retest session, the RTs became shorter compared to the 2 hours retest session (*p* < .001). No main effect of Group was detected, *F*_(1,29)_ = 0.78, *p* = .38, η*_p_^2^*= .03, suggesting that the lack of statistically significant RT difference between groups. We did not find group difference in the pattern of change in average RTs throughout the sessions either (revealed by a non-significant Session × Group interaction: *F*_3,87_ = 0.45, *p* = .72, η*_p_^2^* = .02).

The main effect of the Triplet factor was significant *F*_1,29_ = 190.27, *p* < .001, η*_p_^2^* = .87, indicating that the RTs for the high-probability triplets were faster than for the low-probability triplets. The interaction between the Triplet and Group did not reach significance *F*_1,29_ = 0.75, *p* = .39, η*_p_^2^* = .03. However, the interaction of the Triplet and Session factor was significant *F*_3,87_ = 6.52, *p* < .001, η*_p_^2^* = .18, indicating that the RTs for high- and low-probability triplets changed diversely on the different levels of Session. The pairwise comparisons showed that for the high-probability triplets, the RTs did change between each session except between the 10 minutes and 24 hours retest sessions (*p* = .17, all other *p* < .01). For the low-probability triplets, the RTs did not change between the Training/rTMS session and 2 hours retest session (*p* > .99) and between the 10 minutes retest session and 24 hours retest session (*p* = .59). Triplet learning was influenced by the stimulation as revealed by a significant Session × Triplet × Group interaction *F*_4,116_ = 4.19, *p* = .008, η*_p_^2^* = .13. The pairwise comparisons showed that the RTs for the high- and low-probability triplets did differ in both groups, at each time points (all *p* < .001), and that RTs did not differ between groups in either session either for high- (all *p* > .31) or low-probability triplets (all *p* > .25). This indicates that the difference in the dynamics of the learning curve cannot be explained by solely the changes for the high- or low-probability triplets separately. The subsequent two-way ANOVAs revealed that there was a difference between groups at the 24 hours retest session, Triplet × Group: F_1,29_ = 5.73, *p* = .02, η*_p_^2^* = .17, but not at the initial learning session (*p* = .86), the 10 minutes retest session (*p* = .25) or at the 2 hours retest session (*p* = . 63) (Supplementary Figure 1).

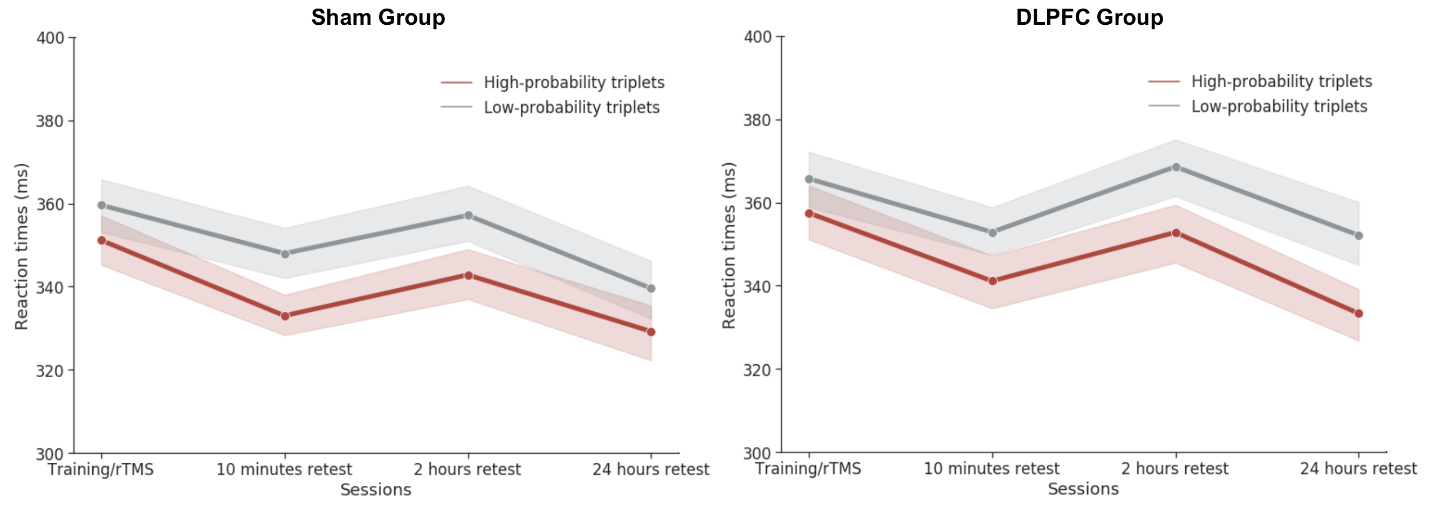

**Supplementary Figure 1. The learning performance in the four experimental sessions.** The vertical axis indicates the RTs for high- and low-probability triplets, while the horizontal axis presents the four experimental sessions. The red line indicates the RT’s for the high-probability triplets, while the gray line the RTs for the low-probability ones. The error bars denote SEM.

**Analysis of the Training/rTMS session**

1) **Learning index**

We also performed an analysis of the Training/rTMS session with the percentage scores. We ran a mixed-design ANOVA with the within-subject factor of Epoch (Epoch 1-5), and the between-subject factor of Group (DLPFC vs. Sham). The main effect of Epoch was found to be significant, *F*_4,116_ = 9.47, *p* < .001, η*_p_*^2^ = .25. Compared to the first epoch, the degree of learning did not change for the second epoch (*p* = .73); however, in the fourth and the fifth epoch, there were significantly higher learning (first epoch vs. fourth epoch: *p* = .001, first epoch vs. fifth epoch: *p* < .001). The interaction of the Epoch and the Group factors were not significant, *F*_4,116_ = .41, *p*  = .80, η*_p_*^2^ = .01, suggesting that the stimulation did not modify how the learning indices changes during the initial learning phase. The main effect of Group was also not significant *F*_1,29_ = 0.04, *p*  = .84, η*_p_*^2^ < .001, indicating that the overall learning in the initial learning indices in this session was not affected by the stimulation (Supplementary Figure 2).

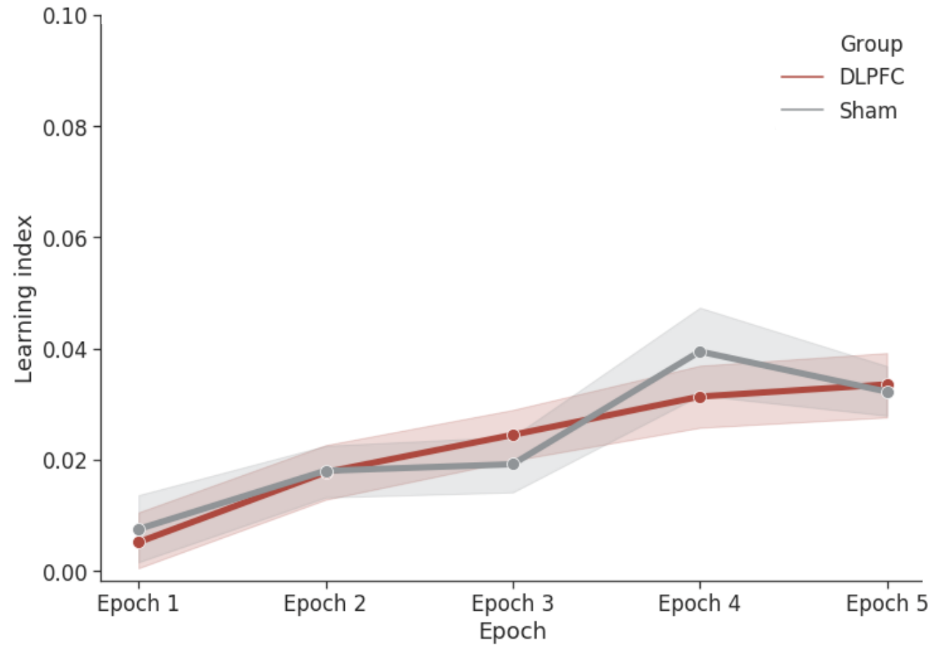

**Supplementary Figure 2. The learning indices of Epoch 1 – Epoch 5 in Session 1.** The vertical axis indicates the learning indices calculated as [(RTs for low-probability triplet – RTs for high-probability triplets)/RT’s for low-probability triplets]. The horizontal axis presents the five epochs of the Training/rTMS session. The red line indicates the learning indices for the DLPFC Group, while the grey line for the Sham Group. The error bars denote SEM.

2) **Raw RTs**

We performed a similar analysis with the raw RT scores for the high- and low-probability triplets. We ran a mixed-design ANOVA with the within-subject factors of Epoch (Epoch 1-5) and Triplet (high- vs. low-probability) and with the between-subject factor of Group (DLPFC Stimulation vs. Sham Stimulation). The main effect of Epoch was significant, F_4,116_ = 86.10, *p* < .001, η*_p_*^2^ = .75. As here the raw RTs were used as dependent variables, the main effect of Epoch indicates that there was a change in the average RTs throughout the five epochs. The pairwise comparisons showed that there was a gradual speed-up of RTs throughout the five epochs. The RTs in the second epoch was significantly shorter than in the first epoch (*p* < .001), the third epoch’s RTs were shorter than the second epoch’s RTs (*p* = .003), and the RTs in the fifth was shorter than in the fourth epoch (*p* = .03). The RTs between the third and the fourth epoch did not change significantly (*p* = .99). No significant main effect of Group was detected *F*_1,29_ = 0.45, *p* = .51, η*_p_*^2^ = .02, suggesting that the average RTs did not differ in the two groups. The Epoch × Group was not significant, *F*_4,116_ = 0.14, *p* = .97, η*_p_*^2^ = .01, revealing that the pattern of change in RTs was not affected by the stimulation.

The main effect of Triplet was significant, *F*_1,29_ = 109.88, *p* < .001, η*_p_*^2^ = .79: the RTs for the high-probability triplets were faster than for the low-probability ones. The interaction of the Triplet and Epoch factor was significant, *F*_4,116_ = 8.24, *p* < .001, η*_p_*^2^ = .22. The pairwise comparisons revealed that for the low-probability triplets, there was no change in RTs between the third and the fourth epoch (*p* > .99) and only a trend was revealed between the second and the third (*p* = .07) and the fourth and the fifth (*p* = .09). In contrast, a gradual change was found between almost every consecutive epoch for the high-probability triplets (except between the fourth and the fifth epoch, *p* =.45, and only a trend was revealed between the third and the fourth epoch, *p* = .06). The difference between the high- and low-probability triplets did not reach significance in the first epoch (*p* = .11), but it did for the last four epochs (all *p* < .001). The interaction between the Triplet and Group did not reach significance, *F*_1,29_ = 0.02, *p* = .89, η*_p_*^2^ = .001, suggesting that, overall, the degree of learning was not affected by the stimulation. Moreover, the Epoch × Triplet × Group interaction was not significant either, *F*_4,116_ = 0.40, *p* = .81, η*_p_*^2^ = .01, suggesting that the change in the difference between the high- and low-probability triplets was not affected by the stimulation (Supplementary Figure 3).

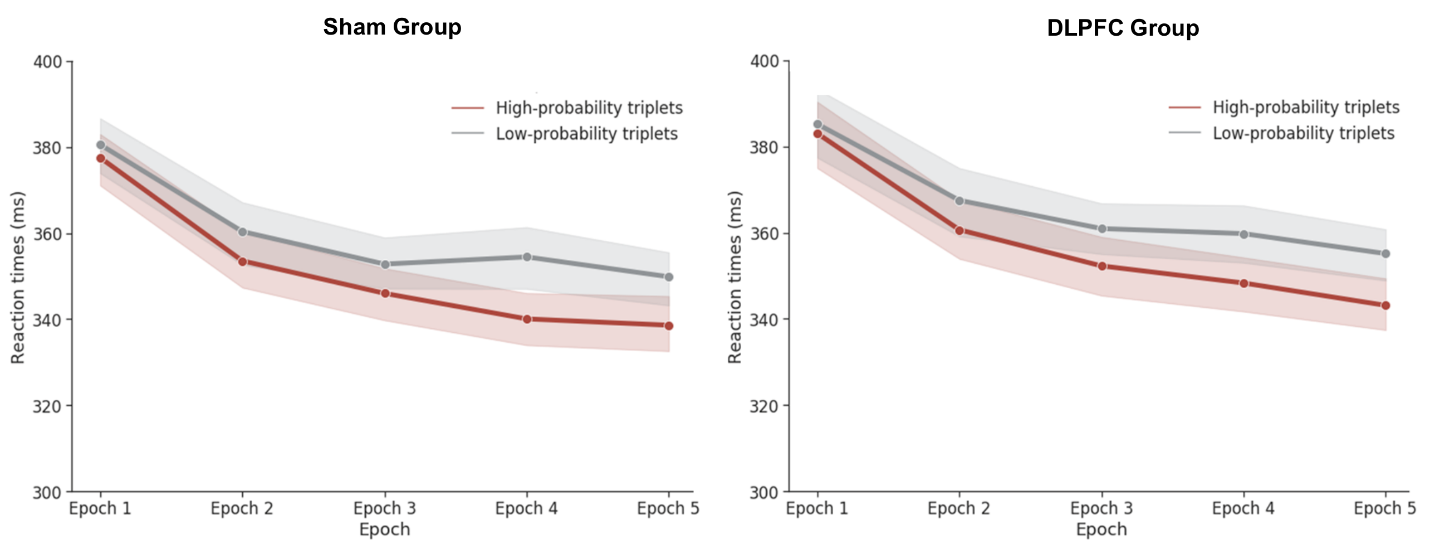

**Supplementary Figure 3. The raw RTs of Epoch 1 – Epoch 5 in Session 1.** The vertical axis indicates the RTs, while the horizontal axis presents the five epochs of the Training/rTMS session. The red line indicates the RT’s for the high-probability triplets, while the gray line the RTs for the low-probability ones. The error bars denote SEM.

**Comparing the performance in the last epoch of the Training/rTMS session to the subsequent sessions**

To check how the performance changed after stimulation has worn off, we performed an analysis of the last epoch of the Training/rTMS session and three subsequent retest session. A mixed-design ANOVA with the within-subject factor of Session (5^th^ Epoch of the Training/rTMS session vs. 10 minutes retest session vs. 2 hours retest session vs. 24 hours retest Session) and the between-subject factor of Group (DLPFC Stimulation vs. Sham Stimulation) was run on the percentage learning indices as described above. The main effect of Session was not significant, *F*_3,87_ = 1.22, *p* = .31, η*_p_*^2^ = .04, indicating that, overall, the degree of learning did not change from the end of the Training/rTMS session. The main effect of Group was also not significant, *F*_1,29_ = .66, *p*  = .42, η*_p_*^2^ = .02, suggesting that the overall learning score was not affected by the stimulation. Importantly, the interaction of the Session and Group factors was significant, *F*_3,87_ = 3.29, *p*  = .03, η*_p_^2^* = .10, indicating that the degree of learning in the four sessions was differently affected by the stimulation. The pairwise comparisons showed that in the Sham Group, the learning indices did not change across the sessions (all *p* > .60). In contrast, in the DLPFC Group, the learning indices showed a trend-level improvement between the Training/rTMS session and the 24 hours retest session (*p* = .10), and between the 10 minutes retest session and the 24 hours retest session (*p* = .08). Moreover, learning was higher for the DLPFC Group at 24 hours retest session compared to the Sham Group (*p* = .03), but not at the other three sessions (all *p* > .25) (Supplementary Figure 4).

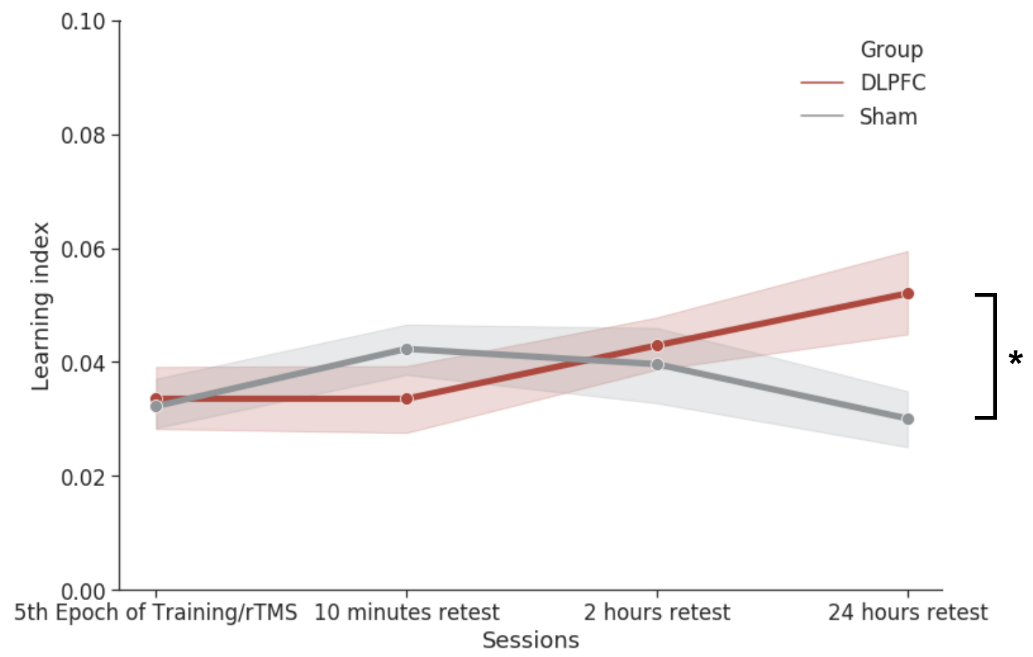

**Supplementary Figure 4. The learning indices of the four experimental sessions.** The vertical axis indicates the learning indices calculated as [(RTs for low-probability triplet – RTs for high-probability triplets)/RT’s for low-probability triplets]. The horizontal axis presents the four experimental sessions (for the Training/rTMS session, only the last epoch is included). The red line indicates the learning indices for the DLPFC Group, while the grey line for the Sham Group. The error bars denote SEM.
